# Opposite-sex pairing alters social interaction-induced GCaMP and dopamine activity in the insular cortex of male prairie voles

**DOI:** 10.1101/2024.11.21.624717

**Authors:** Erika M. Vitale, Amina H. Tbaba, Kaitlyn Tam, Kyle R. Gossman, Adam S. Smith

## Abstract

The prairie vole (*Microtus ochrogaster*) is a monogamous rodent species which displays selective social behaviors to conspecifics after establishing a pair bonded relationship, specifically partner-directed affiliation and stranger-directed aggression. This social selectivity relies on the ability of an individual to respond appropriately to a social context and requires salience detection and valence assignment. The anterior insular cortex (aIC) has been implicated in stimulus processing and categorization across a variety of contexts and is well-situated to integrate environmental stimuli and internal affective states to modulate complex goal-directed behaviors and social decision-making. Surprisingly, the contribution of the aIC to the expression of pair bond-induced social selectivity in prairie voles has been drastically understudied. Here we examined whether neural activity and gene expression in the aIC change in response to opposite-sex pairing and/or as a function of pairing length in male prairie voles. Opposite-sex pairing was characterized by changes to calcium and dopamine (DA) transients in the aIC that corresponded with the display of social selectivity across pair bond maturation. Furthermore, D1 and D2 receptor mRNA expression was significantly higher in males after 48 hrs of cohabitation with a female partner compared to same-sex housed males, and D2 mRNA remained significantly higher in males with a female partner compared to same-sex housed males after a week of cohabitation. Together, these results implicate a role for DA and its receptors in the aIC across the transition from early-to late-phase pair bonding.

---

For virtually all social beings, ranging from insect to primate species, navigating a social environment is commonplace and often necessary for both individual and species survival. Effective social communication relies on one’s ability to integrate and process multiple social and environmental modalities to drive an appropriate behavioral response [1, 2]. Examples of such modalities include the external environmental context, the internal emotional and motivational state of the individual, and the perceived internal state(s) of other social conspecifics [3]. Furthermore, social encounters are dyadic and often involve in-the-moment adaptations to the behavioral responses of social partners throughout the length of an interaction session. Social neuroscience has uncovered a network of brain regions that work together to attune, process, and respond appropriately to specific social encounters, and this collection of regions is known as the social decision-making network [4, 5]. The insular cortex (IC) and other “higher order” cortical regions have functional-structural relationships to regions of the social decision-making network but have received less attention regarding its regulation of social behavior [6]. Structural connectivity patterns implicate the posterior IC (pIC) as the main sensory “detector” and relays this information directly to the amygdala for immediate survival-dependent action [7-9]. In contrast, the anterior IC (aIC) receives cortical information, interoceptive information from the hindbrain, and sensory information relayed from the pIC and integrates these signals together to create a wholistic framework for a specific context, then “superimposes” this framework over the motivation-centered striatum to adjust stimuli valence and salience for more nuanced control of reward-based decision-making [10]. Thus, it is no surprise that the IC has been implicated in a wide range of context-dependent learning, memory, and decision-making such as drug and alcohol abuse, taste recognition memory, and social recognition memory [11-16].

Given the structural and functional intricacy of the IC and the fact that activity of the IC has been extensively linked to the expression of a variety of human attachments, including mother-infant bonds and adult romantic relationships [17], it is surprising that the IC has been largely unexplored in basic research using model animal species that form similar attachments. The prairie vole (*Microtus ochrogaster*) is a socially monogamous rodent species that is often used to study the neurobiology of adult social attachments [18, 19]. While researchers have spent decades exploring the neurochemicals and circuits that underly the formation and maintenance of such attachments in prairie voles (termed pair bonds), no studies have focused on the IC. Furthermore, the mesolimbic reward system, both at the level of the cell bodies in the ventral tegmental area (VTA) [20, 21] and downstream in the ventral striatum [22], is crucial for pair bonding. The aIC would be well situated to modulate the behavioral expression of pair-bonding, characterized by partner-specific affiliation and stranger-specific aggression, by receiving contextual information from the pIC and relaying it to the ventral striatum and influence social decision-making.

We hypothesized that the aIC would be differentially recruited during social encounters in bachelor vs pair bonded prairie voles. Furthermore, we predicted the pattern of aIC activity in pair bonded voles would vary based on the length of pairing (short-term vs long-term) and social stimulus type (familiar vs unfamiliar). We used fiber photometry to assess calcium and DA transients in the aIC during partner and stranger encounters. To determine whether pair bonding is associated with post-synaptic changes in the aIC, we used qRT-PCR to analyze mRNA expression of DA receptor types 1 (DRD1) and 2 (DRD2) in short-term and long-term pair bonded prairie voles.

## Methods

### Animals

Prairie voles were lab-bred from a population captured in southern Illinois. Voles were weaned at 21 ± 3 days of age and were housed in same-sex, non-sibling pairs in microisolator cages (29.2L x 19.1W x 12.7H cm) with corn cob bedding, crinkle nesting material, and ad libitum access to food (Tekland global rabbit diet 2030) and water. Colony rooms were maintained at 21 ±1°C with a 14L:10D photoperiod (lights on at 0600 h). Male subjects were between 90 and 120 days of age at the start of the experiment. Female prairie voles (also between 90 and 120 days of age) were used as partners and had been tubal ligated at least one week before pairing (see [23] for description of tubal ligation procedure). All behavior testing was performed between 0900-1700h, and all procedures were conducted in accordance with the National Institutes of Health Guide for the Care and Use of Laboratory Animals and the Institutional Animal Care and Use Committee at the University of Kansas.

### Experiment 1: Effects of same-vs opposite-sex cohabitation on social stimuli-induced aIC activity

#### Experimental Design

Sixteen male prairie voles were used as subjects (n = 8/group). Subjects received stereotaxic surgery (see below for surgery details) then remained housed with their same-sex, non-sibling cage mate throughout the recovery and viral incubation period of three weeks. For subjects in the same-sex (SS) paired group, only one animal from the cage received surgery and they recovered and remained with their same-sex cage mate throughout the entirety of the experiment. Subjects in the opposite-sex (OS) paired group were housed with a non-related, tubal ligated female immediately following the social exposure test on Pairing Day (Experimental Day 0).

#### Stereotaxic Surgery

Subjects were anesthetized with an intraperitoneal injection of a ketamine (75mg/kg) and dexmedetomidine (1mg/kg) cocktail. The head was shaved, and the skin was cleaned with 70% ethanol then betadine solutions and repeated three times. A subcutaneous injection of a local anesthetic (lidocaine, 2mg/kg) was administered at the surgical site. The head was fixed into a stereotaxic apparatus (Stoelting), and an incision was made exposing the skull. A hole was drilled above the injection site, and a 1 uL Neuros needle (Hamilton) was slowly lowered to the rostral insular cortex (AP +1.60 mm, ML +3.25 mm, DV -4.00 mm from bregma). A viral cocktail consisting of 150 nL of pAAV1-hSyn-GCaMP6f-WPRE-SV40 (AddGene) and 150 nL of pAAV9-hSyn-GRAB-rDA1m (AddGene) was infused at a rate of 60 nL/min followed by a 5 min diffusion period. Two surgical screws were fixed to the skull, and a fiber optic implant (1.25mm ceramic ferrule, 200μm core, 0.5NA, 6mm length; RWD) was lowered to the same aIC coordinates with the DV adjusted to -3.80 mm. Dental cement was used to secure the implant and screws, then antisedan (2.5mg/kg) was administered to reverse the anesthesia. Subjects received 2 doses of meloxicam (2mg/kg) post-operation (over a 24-hour period) and were also permanently re-housed with their cage mate in a slightly larger cage (36.2L x 20.3W x 14.0H cm) with more vertical clearance from the hopper to accommodate the cranial implant. Animals remained unmanipulated aside from regular facility cage changes for 3 weeks to allow for complete viral transfection.

#### Fiber Photometry & Social Exposure Tests

On the first day of experimental testing (Pairing Day = Experimental Day 0), subjects were brought to the behavioral testing suite and allowed to habituate for 1 hour. Next, a subject was removed from the home cage, the optical implant was attached to a fiber optic cable (Plexon) via a ceramic ferrule, and the animal was placed into a clean, novel cage (47.6L x 26.0W x 15.2H cm) and allowed to habituate for 10 min. A baseline (no behavior) recording was captured via a Plexon Multi-Channel photometry system for 5 min before two 20-min social exposure test sessions (i.e. Session A and Session B): SS males interacting with their same-sex cagemate or a novel same-sex conspecific and OS males interacting with two different novel females (one which will serve as their partner moving forward). Stimulus order was randomized between subjects, and there was a 20 min nonsocial “rest” period between each social exposure. Immediately following Session B, males in the OS group were permanently housed with one of the two females they had interacted with during the social exposure test. Social exposure tests were also conducted on Experimental days 2 and 8, with all partner interactions occurring with the same partner/cage mate while all stranger interactions occurred with a conspecific that the subject had never previously interacted with. All social exposure behavior sessions were video recorded at 30 frames per second using a Logitech web camera. Fiber photometry data from the 410, 450, and 560 nm channels were captured at 30 fps. Cameras were connected to an input relay device attached to the Plexon Multi-Channel photometry system that signaled the start of the video recording to the system for time-locking behavioral assessments with the photometry signals. Social behaviors were coded frame-by-frame using Solomon Coder, and timestamps for the onset of all behaviors were extracted and used as events for photometry analysis. Social behaviors were analyzed for frequency and duration, including social proximity, olfactory investigation, affiliative behaviors (side-by-side, huddling, allogrooming), and agonistic behaviors (boxing, defensive treading, fleeing from the stimulus, and chase/tumble sequences).

#### Tissue Collection and Confirmation of Virus Placement

At the end of the experiment, subjects were deeply anesthetized with a ketamine/dexmedetomidine cocktail, then perfused with saline and 4% paraformaldehyde in 0.1M Phosphate Buffer solution (pH = 7.4). Brains were extracted, post-fixed overnight, transferred to 30% sucrose, and sectioned at 30 μm. Every other section through the aIC/mIC (+1.90mm-0.20mm from bregma) was mounted onto glass slides and coverslipped using a hard-set, antifade coverslipping medium (Gelvatol, made in-house). Once dry, sections were viewed under a Leica microscope using a chroma 488 filter (L5-ET, Leica) to visualize the GFP-conjugated GCaMP virus and a 555 nm filter (RHOD-ET, Leica) to visualize the mApple-conjugated GRAB_DA_. Animals were removed from the study if viral expression or optical fiber implant was found outside of the IC.

#### Fiber Photometry Analysis

Raw fiber photometry output for the 410, 450, and 560 nm channels was exported from Plexon Software, time-locked to the start of video recording, then analyzed using open-source fiber photometry analysis software (Guided Photometry Analysis in Python; GuPPy) [24]. Motion artifact was corrected for in GuPPy by using raw signal from the isobestic (410) channel. Timestamps for each behavior were used as events to examine event-related changes in aIC calcium and DA activity corresponding to specific social behaviors. Area under the curve (AUC) was calculated for both anticipatory (−2 to 0 sec prior to behavior onset) and initiation-induced (0 to 5 sec) brain activity responses to each social behavior.

### Experiment 2: Effects of same-vs opposite-sex cohabitation on DA receptor mRNA expression

#### Experimental Design, Brain Collection, and qRT-PCR

A separate cohort of male prairie voles (N=36) were divided into same-sex paired (SS, n=11), and opposite-sex paired groups that cohabitated with a tubal ligated female for either 48 h (OS-ST, n=13) or 1 week (OS-LT, n=12). Immediately prior to tissue collection, subjects were tested for the expression of a pair bond using the partner preference test (PPT; pair bonding was characterized by an animal spending greater than or equal to a 3:1 ratio of affiliation with their partner vs the stranger, [25]). Subjects were rapidly decapitated, and brains were extracted and stored at -80°C before being sectioned at 200 μm using a cryostat (Leica). Unilateral tissue punches (1.0 mm diameter) were collected from the aIC. Tissue punches were homogenized, and RNA was extracted and purified using a Qiagen RNEasy mini kit following manufacturer’s instructions. RNA was quantified using a Qubit 3 fluorometer and a high sensitivity RNA quantification kit (Thermo Fisher Scientific, Waltham, MA). 20-100 ng of mRNA was converted to cDNA using a high-capacity reverse transcription kit (Applied Biosystems, Foster City, CA) per the manufacturer’s instructions. mRNA for dopamine receptor 1 (DRD1) and dopamine receptor 2 (DRD2) were analyzed using probes designed from target gene sequences of the prairie vole genome [25]. Hypoxanthine-guanine phosphoribosyltransferase (HPRT) was used as the comparison “housekeeping” gene based on its relatively constant expression in cells independent of experimental conditions [26, 27]. qRT-PCR for each target was run in triplicate for every subject on one plate with wells containing 5 ng cDNA, SYBR green PCR Master Mix (Applied Biosystems, Foster City, CA) and 200 nM of each forward and reverse primer (see [25] for specific sequences). A ThermoFisher StepOnePlus PCR plate reader was used for quantification. A dissociation curve was generated for each sample and used to confirm that only a single product was transcribed. The ΔΔCT method was used to calculate the fold differences between groups [28].

#### Data Analyses

Significant differences were determined by a p-value of <0.05 for all analyses. In Experiment 1, social behaviors and behavior-related AUC results were analyzed using linear mixed modeling (LMM) using test session (Pairing Day, ST, LT) and stimulus type (Partner vs Stranger) as within-groups variables and companion sex (SS vs OS paired) as a between groups variable. Significant main effects and interactions were followed by Bonferroni post-hoc analyses to further elucidate the data patterns. The mRNA results in Experiment 2 violated the One-Way ANOVA assumption of homogeneity of variances between groups, so Kruskal-Wallis was used to analyze group differences in DRD1 and DRD2 expression. Significant results were followed by Bonferroni post-hoc analyses.

## Results

### Experiment 1-1: Companion sex and pairing length influence partner- and stranger-directed social behaviors

All subjects spent more time in social proximity (F=8.234, p = 0.008) and side-by-side contact (F = 7.19, p = 0.013) with their partner compared to the stranger conspecific and were more aggressive to the stranger than partner (F = 7.19, p = 0.013). However, there were significant 3-way interactions between time, companion sex, and social stimulus type for each of these behaviors. Specifically, SS males spent significantly more time in proximity (Fig 2A) and side-by-side contact (Fig 4A) with their partner compared to a stranger on the first day of testing but not subsequent test periods. In contrast, OS males were in proximity (Fig 2B) and side-by-side contact (Fig 4A) to the two novel females during the first encounter, but males spent more time in proximity and side-by-side contact to their female partner than the stranger female after cohabitating with their partner for two and seven days. In addition, SS males showed the most stranger-directed aggression during the first social exposure test while OS males were not aggressive before cohabitation then became significantly more aggressive towards the stranger during subsequent social exposure tests (ST and LT timepoints).

**Figure 1.**
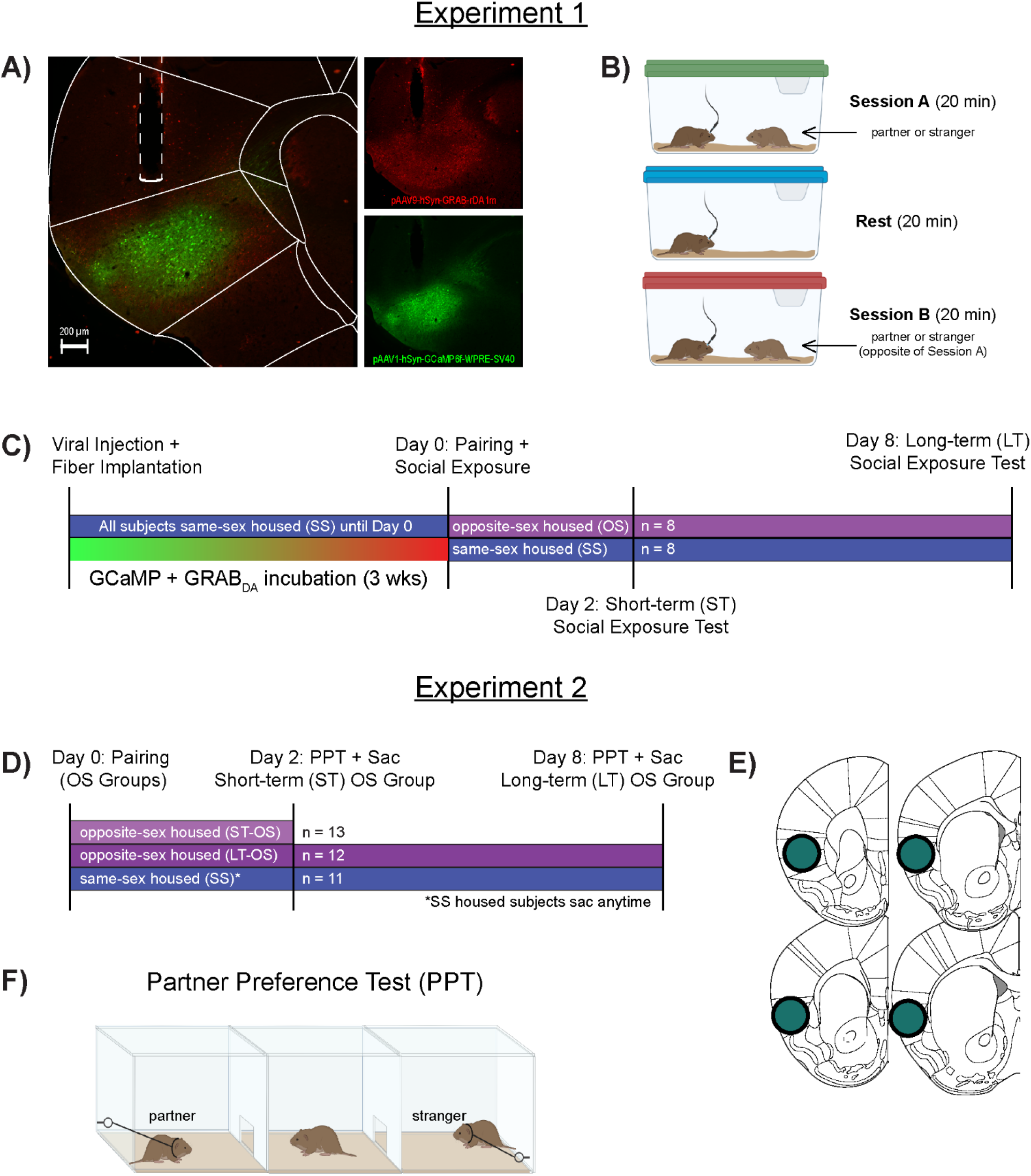
Schematic of experimental procedures and timelines. (A) GRAB_DA_ and GCaMP expression and optical fiber placement in the anterior insular cortex. (B) Fiber Photometry recording was performed during each social exposure session. Subjects were exposed to one of two potential social conspecifics during Session A (partner or stranger), followed by a non-social “rest” period. During Session B, subjects were exposed to whichever social stimuli was not introduced in Session A. (C) Viral injection and fiber implantation occurred 3 weeks before the start of the experiment. On Day 0, subjects were randomly assigned to either the same-sex (SS) or opposite-sex (OS) paired groups. SS paired males remained with their same-sex cage mate and were exposed to their cage mate and a stranger during the Day 0 social exposure test. OS paired males were exposed to two novel females during the Day 0 social exposure, with one of these females becoming the subjects’ partner immediately following the test. Social exposure tests also occurred on Days 2 (ST pairing) and 8 (LT pairing). (D) A separate cohort of male subjects were randomly divided into 3 groups: SS housed subjects were never paired with a female, while two OS groups were cohabitated with a female for either 2 days (ST-OS) or 1 week (LT-OS). Brains were collected from all three groups and tissue was used for qRT-PCR analysis. (E) Tissue punches were collected unilaterally through the anterior IC. (F) A partner preference test was used to confirm pair bonding in both OS groups and occurred several hours before subjects were sacrificed.

**Figure 2.**
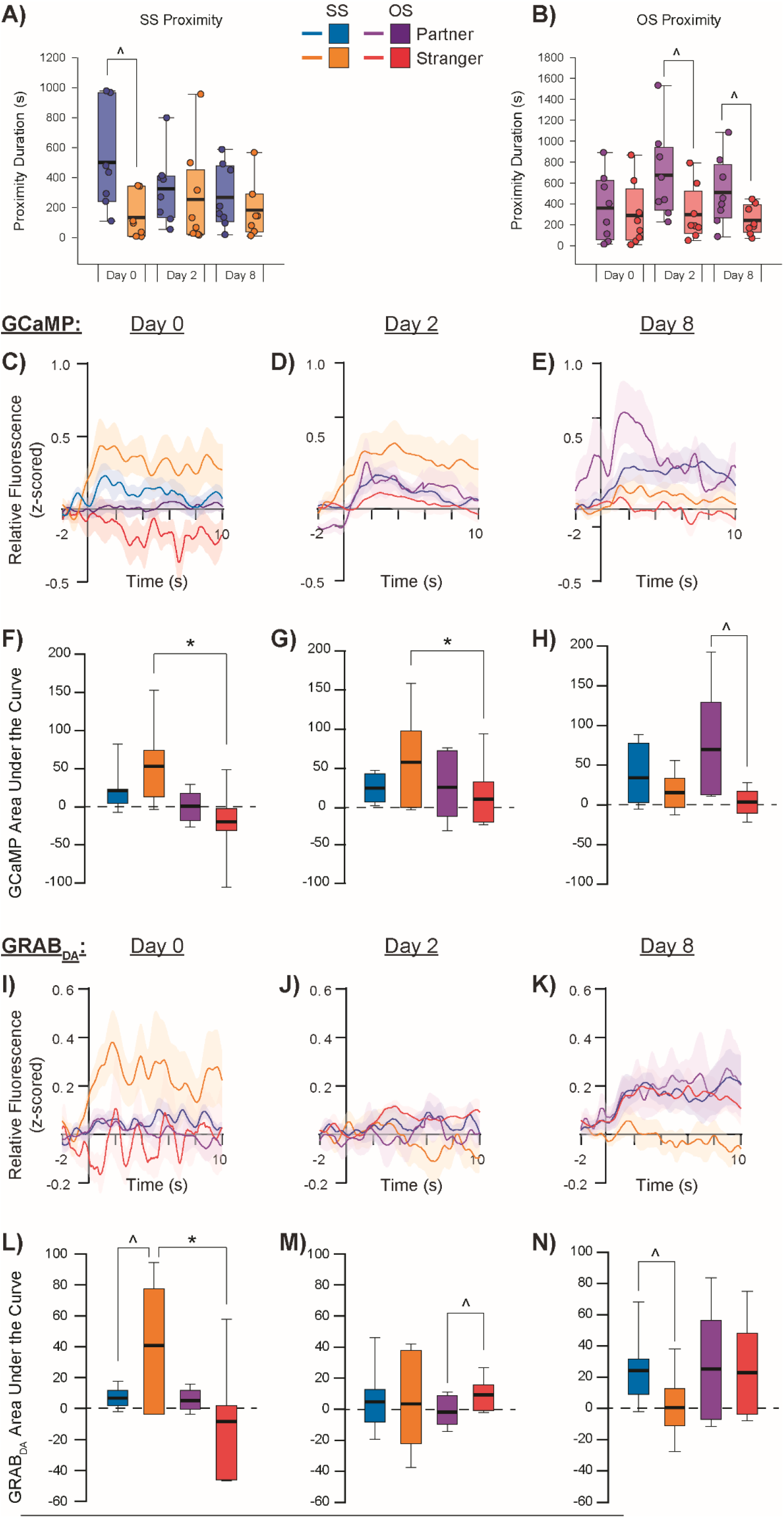
Social proximity-induced GCaMP and GRAB_DA_ activity in the aIC. SS paired subjects (A) and OS paired subjects (B) show differing patterns of close proximity to partners vs strangers across testing days. GCaMP relative fluorescence during close proximity to partners vs strangers in SS and OS paired subjects during social exposure tests on experimental Day 0 (C), Day 2 (D), and Day 8 (E) with corresponding area under the curve (AUC) calculations (F-H). GRAB_DA_ relative fluorescence during close proximity to partners vs strangers in SS and OS paired subjects during social exposure tests on Day 0 (I), Day 2 (J), and Day 8 (K) with corresponding AUC calculations (L-N). ^denotes p<0.05 for partner vs stranger within pairing group, *denotes p<0.05 for SS vs OS pairing condition within the same social stimulus type.

### Experiment 1-2: Companion sex and pairing length influences social behavior-related GCaMP and GRAB_*DA*_ signal in the aIC

There were several behavior-dependent main effects and interactions for GCaMP and GRAB_DA_ responses across different conditions. When subjects initiated proximity with a social stimulus, there was a significant difference in the size (AUC) of GCaMP transients between SS and OS subjects, with GCaMP in the aIC showing a larger proximity-induced response overall in SS subjects compared to OS subjects (F = 5.170, p = 0.026). However, this main effect of companion sex appears to be driven by an interaction with stimulus type (F = 8.852, p = 0.004), with SS males showing a substantially higher total aIC GCaMP response to proximity with strangers compared to OS males in proximity to strangers (Bonferroni corrected pairwise comparison t = 3.695, p < 001; Fig 2C-H). There was also a significant interaction between companion sex and test day, such that SS males showed a larger response than OS males on the first (pairing; t = 3.973, p < 0.001) test day but not the second (short-term paired) or the final (long-term paired) test days. Finally, there was a significant interaction between social stimulus type and testing day (F = 9.206; p < 0.001), with there being a significantly higher response to proximity with the partner compared to the stranger on the final test day (regardless of SS or OS pairing; t = 3.954, p < 0.001). aIC GCaMP response to partner proximity in OS males increased significantly as pairing length continued, while this pattern did not emerge for OS males in proximity to strangers or SS males in proximity to either their partner or to strangers.

GRAB_DA_ transient size (AUC) in the aIC during social proximity changed significantly across testing day (F = 11.133, p < 0.001). However, this main effect was driven by a significant 3-way interaction between companion sex, stimulus type, and test day (F = 5.761, p = 0.005; Fig 2I-N). Specifically, SS males showed a greater DA response to stranger proximity than OS males on the first testing day (t = 2.186, p = 0.032). Furthermore, this DA response to stranger proximity in SS males also significantly differed from their DA response to partner proximity, with an inversion of response occurring where strangers elicited significantly greater DA release on the first test day (t = 2.019, p = 0.047) whereas partners elicited greater DA release on the last test day (t = 3.507, p < 0.001). DA activity in the aIC of OS males in social proximity did not differ across social stimulus type or testing day.

During social olfactory investigation, SS males showed a significantly greater overall GCaMP response compared to OS males (F = 6.882, p = 0.011; Fig 3C-H). DA response, however, showed a significant interaction between companion sex and test day (F = 3.836, p = 0.026), with SS males having an overall larger DA response to olfactory investigation of strangers than OS males did when investigating strangers (t = 2.030, p = 0.046; Fig 3I-N). Finally, DA response to olfactory investigation between partners and strangers in SS males showed the same pattern as proximity, where sniffing strangers elicited greater DA activity on the first test day (t = 2.276, p = 0.026), while sniffing partners elicited more DA activity than sniffing strangers did on the subsequent testing days (short-term: t = 2.034, p = 0.046; long-term: t = 2.271, p = 0.026).

**Figure 3.**
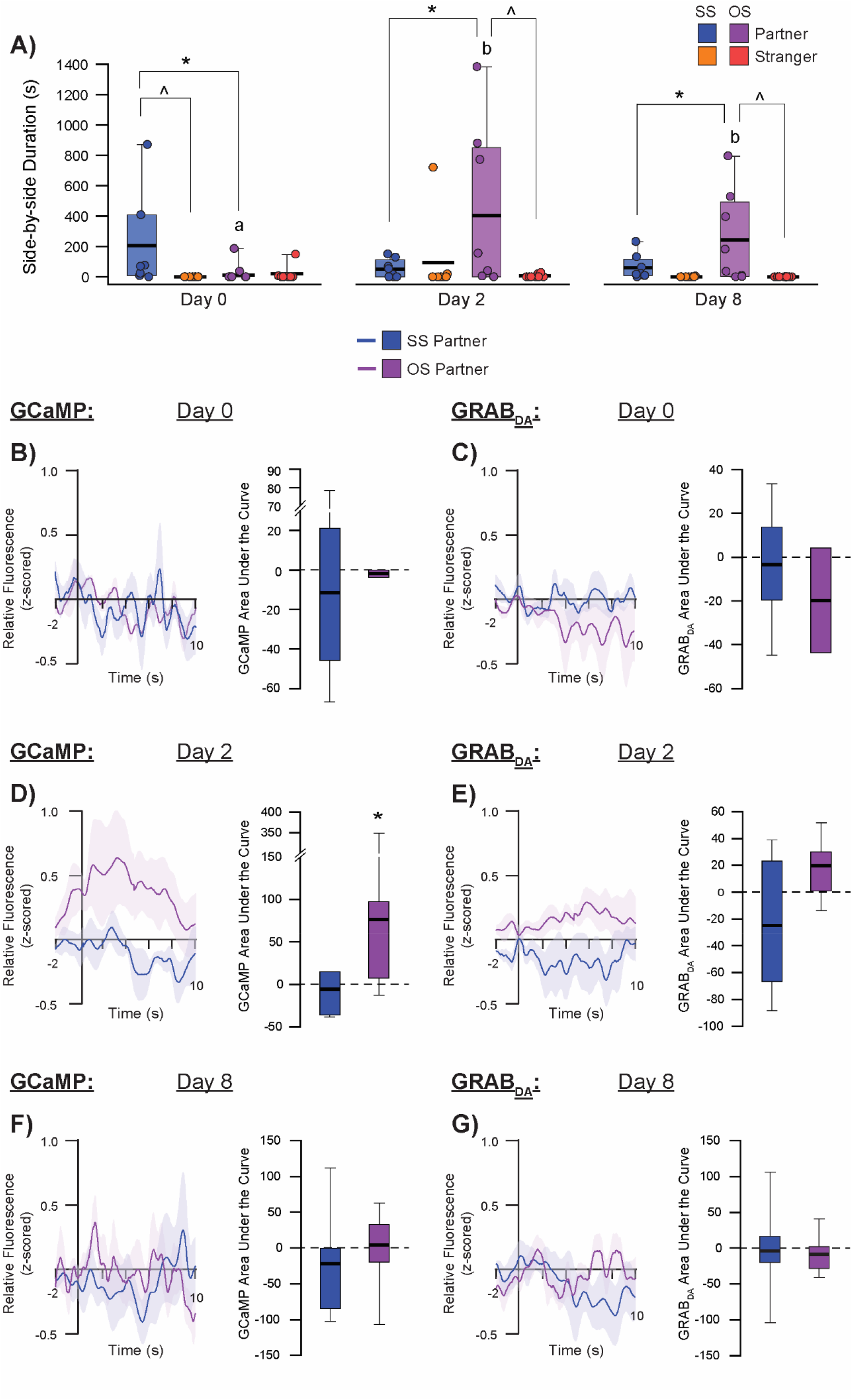
Olfactory investigation-induces GCaMP and GRAB_DA_ activity in the aIC. SS paired subjects (A) and OS paired subjects (B) do not significantly differ in their patterns of olfactory investigation of partners vs strangers across testing days. GCaMP relative fluorescence during olfactory investigation of partners vs strangers in SS and OS paired subjects during social exposure tests on experimental Day 0 (C), Day 2 (D), and Day 8 (E) with corresponding area under the curve (AUC) calculations (F-H). GRAB_DA_ relative fluorescence during olfactory investigation of partners vs strangers in SS and OS paired subjects during social exposure tests on Day 0 (I), Day 2 (J), and Day 8 (K) with corresponding AUC calculations (L-N). ^denotes p<0.05 for partner vs stranger within pairing group, *denotes p<0.05 for SS vs OS pairing condition within the same social stimulus type.

**Figure 4.**
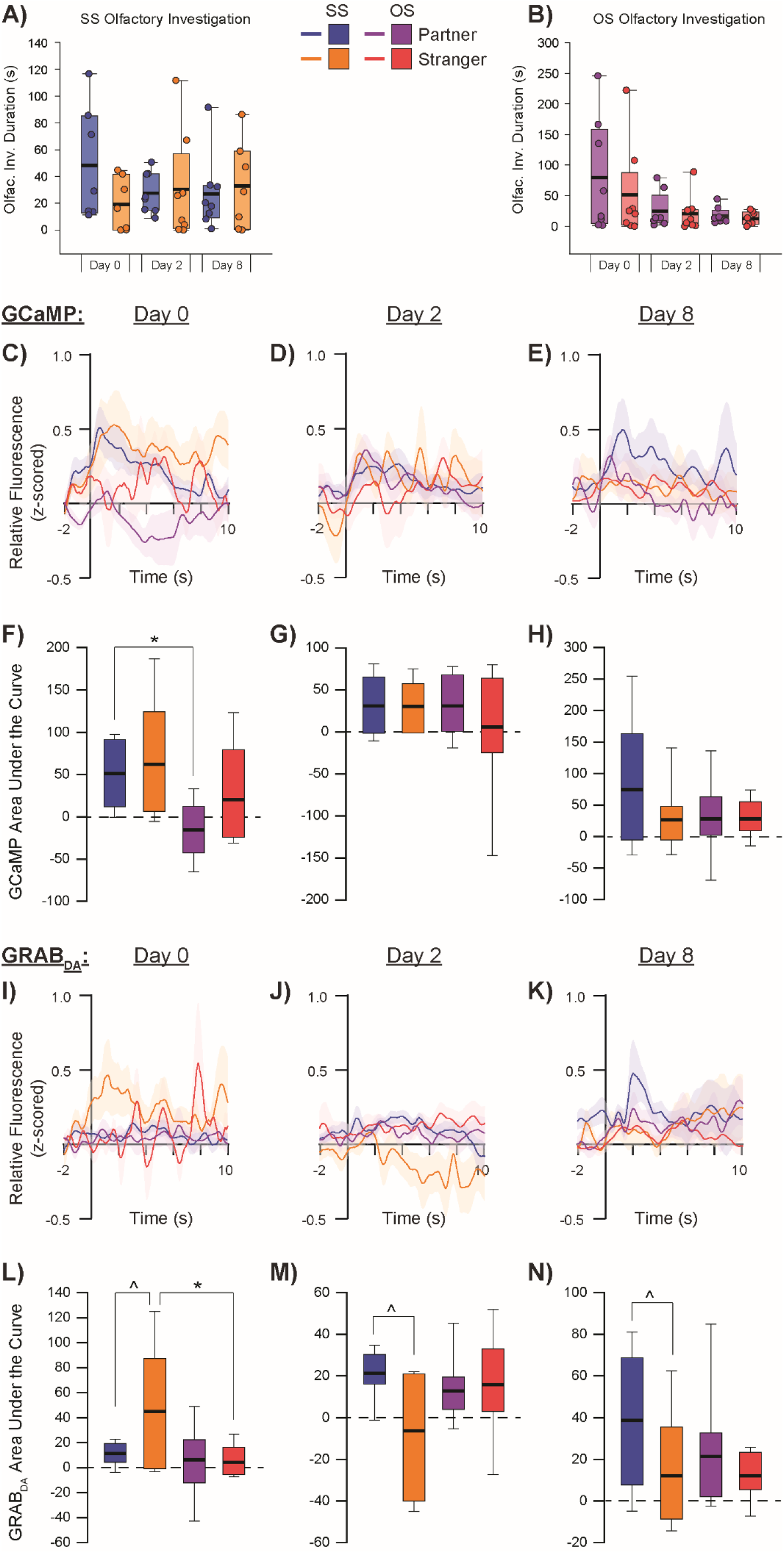
Affiliation-induced GCaMP and GRAB_DA_ activity in the aIC. SS and OS paired subjects show selective affiliation with their partners and rarely affiliate with strangers (A). GCaMP relative fluorescence and AUC (B) and GRAB_DA_ relative fluorescence and AUC (C) during side-by-side contact on Day 0 partner exposure session, Day 2 partner exposure session (D-E), and Day 8 partner exposure session (F-G). ^denotes p<0.05 for partner vs stranger within pairing group, *denotes p<0.05 for SS vs OS pairing condition within the same social stimulus type, different letters denote p<0.05 for same stimulus type across testing sessions for OS paired subjects.

**Figure 5.**
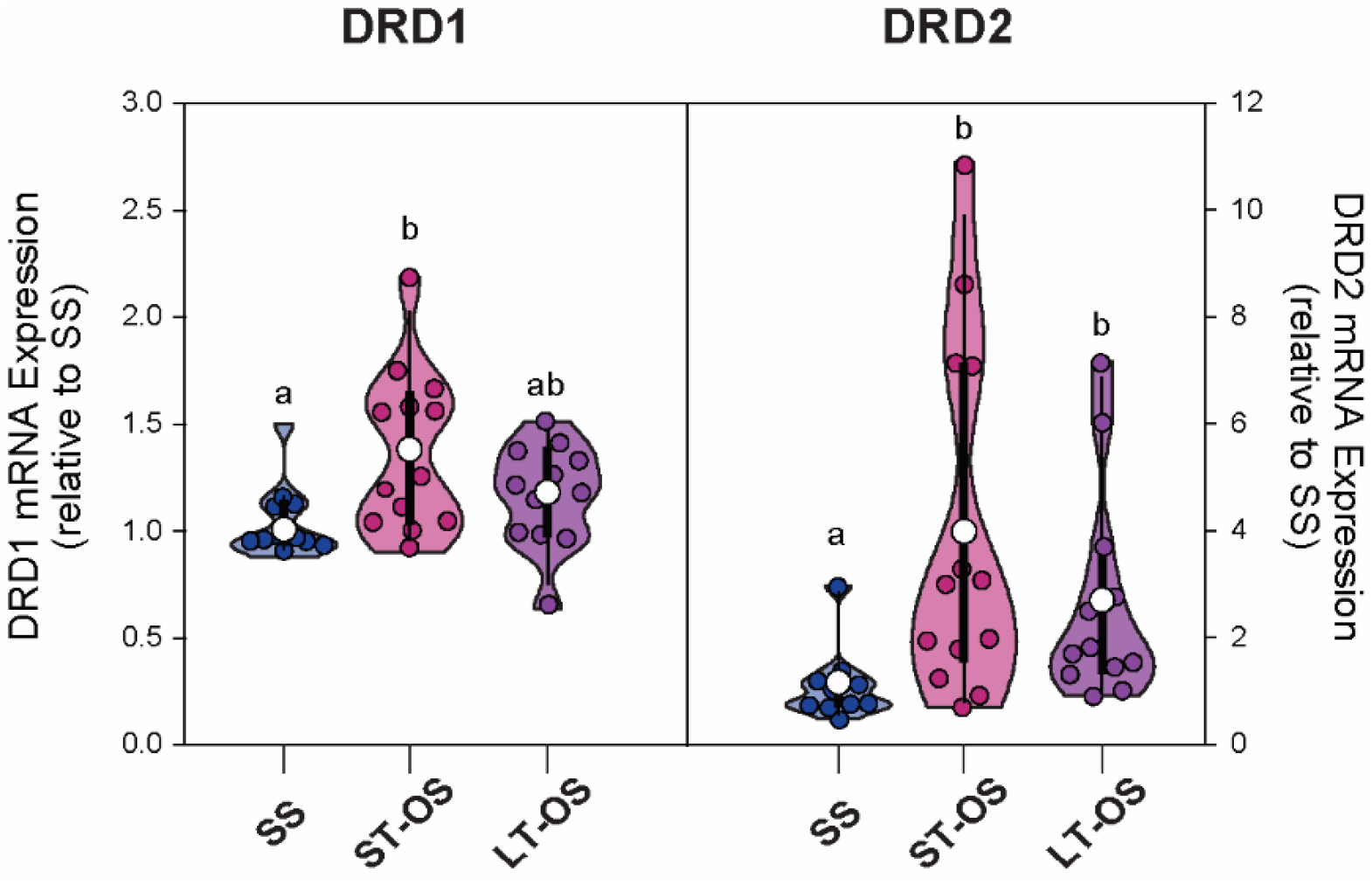
DRD1 and DRD2 mRNA expression in the aIC. OS pairing is characterized by a selective increase DRD1 mRNA after 2 days but not 1 week of opposite-sex cohabitation in comparison to SS paired subjects. DRD2 mRNA expression is significantly higher in both OS pairing groups compared to SS males. Different letters denote p<0.05 between groups.

Upon initiation of side-by-side contact with partners, OS males showed a significantly greater aIC GCaMP response compared to SS males (main effect of companion sex: F = 4.494, p = 0.046). Similarly, there appeared to be a greater anticipatory response in OS males, as indicated by a significantly higher area under the curve during the 2 sec immediately before initiation of side-by-side contact compared to SS males (F = 8.313, p = 0.007). Although there were no significant effects of side-by-side contact initiation with the partner on DA activity in the aIC based on companion sex or testing day, there was an anticipatory interaction between companion sex and testing day (F = 12.203, p < 0.001). Specifically, OS males showed significantly greater DA activity just before (2 sec) initiation of side-by-side contact with their partner on the second day of testing (after 48 hrs of pairing) compared to SS males (t = 3.483, p = 0.002). This DA response to anticipation of side-by-side partner contact in OS males after 48 hours of pairing was also significantly higher than the response on pairing day and after one week of pairing.

There was a significant effect of companion sex on GCaMP activity to initiation of stranger-directed aggression where SS males showed a significantly higher total response compared to OS males (F = 5.468, p = 0.028). There were no effects of companion sex or test day on DA activity in response to stranger-directed aggression, nor were there any anticipatory effects for either GCaMP or DA activity.

### Experiment 2: Opposite-sex cohabitation alters DA receptor mRNA expression in the aIC

There was a significant group effect on mRNA expression of DRD1 (F_(33,2)_ = 6.911, p = 0.032) and DRD2 (F_(32,2)_ = 11.096, p = 0.004) receptors in the aIC. Specifically, subjects in the OS-ST pairing group had significantly higher DRD1 mRNA expression compared to the SS group (t = 11.343, p = 0.026), while both opposite-sex pairing groups had significantly higher DRD2 mRNA expression compared to the SS group in the aIC (OS-ST vs SS: t = 13.90, p = 0.004; OS-LT vs SS: t = 10.90, p = 0.039).

## Discussion

Prairie voles increased display of partner-directed affiliation and stranger-directed aggression as the length of opposite-sex pairing increases, findings that replicate previous work [23, 29]. Furthermore, during social encounters with a novel social stimulus, proximity-induced aIC activity was significantly lower in pair bonded males than bachelor males, most notably during the first and second novel encounter sessions. Bachelor males also had significantly greater GCaMP transients to stranger proximity than partner proximity, suggesting that the bachelor aIC responds more to social novelty. A recent study in mice showed that interacting with a novel conspecific increased calcium activity in most socially-responsive aIC neurons, particularly during stationary contact behavior [30]. This aligns with the proposed role of the aIC in social decision-making, as the bulk of its social interaction-induced calcium transients occurs during stationary “processing” phases of the social encounter then relays the behavioral “decision” to other brain regions [10]. Social-related aIC activity also appears to be modulated by pair bonding, as evidenced by significantly greater aIC GCaMP transients to partner proximity compared to stranger proximity in the pair bonded males. Bachelor males also exhibited stable, positive GCaMP transients during peer proximity throughout the experiment. Continuous interaction with a social companion may strengthen the pathways from sensory-related areas to the IC to facilitate recognition of a familiar social companion. It has recently been shown that excitotoxic lesions to the aIC impair social recognition [12, 31, 32]. Thus, the ability to recognize and associate socio-sensory cues with a particular social partner appears to require activity of the aIC.

DA release and receptor expression have been previously shown to modulate both pair bond formation and maintenance in prairie vole males [33]. For instance, DRD2 receptors in the NAcSh are critical for partner preference formation [22]. Conversely, DRD1 receptor expression is upregulated in the NAcSh after long-term opposite-sex cohabitation, and their activity is required for stranger-directed aggression [34, 35]. DRD1-expressing neurons are also necessary for the display of appetitive aspects of partner motivation since lever pressing for a pair bonded partner causes increased NAcSh DA signaling and blocking DRD1 receptors significantly reduces lever pressing for a partner [36]. Surprisingly, DA-related mechanisms during the transition period from short-term to long-term pair bonding have rarely been studied. Here, we noticed a significant increase in partner-directed side-by-side contact in OS males after 2 days of pairing, and though this behavior remained high after a week of pairing, the average duration did drop slightly. Interestingly, both aIC GCaMP and GRAB_DA_ transients during initiation of side-by-side contact were significantly higher than aIC transients in SS males to side-by-side contact only at this early pairing timepoint. Agmo et al. [37] noted a similar behavioral pattern in pair bonded black tufted-ear marmosets (*Callithrix pencillata*), with huddling and allogrooming behaviors peaking on days 2-6 of pairing and then dropping slightly thereafter. Although this behavioral pattern has not been consistently observed in prairie voles [29], the significant increase in aIC GCaMP activity and DA release during side-by-side contact at this stable but still early phase of a pair bond could signify a neurobiological epoch that forms the foundation of a long-term relationship by linking motivational systems to social contact with their partner. Indeed, both affective touch and physiologic sensory input (or interoception) are relayed from the body to the brain via small, unmyelinated afferents that converge in the mid-posterior IC [38-40], implying that social touch can be a powerful modulator of physiological and emotional state via the IC.

In mice, the activity of DRD1-expressing neurons in the aIC projecting to the lateral NAcSh predicts both affiliative behaviors during a prosocial interaction and agonistic behaviors during an aversive social experience [41]. Thus, social stimuli-dependent synaptic plasticity originating from aIC^D1^ neurons appears to drive contextually appropriate learned social behaviors. Furthermore, the pair bonding process seems to involve its own form of DA-related neural plasticity since DRD1 and DRD2 mRNA expression were increased in the aIC of pair bonded male prairie voles compared to bachelors. This means that while a similar DA response occurs in the aIC during both partner and novel social encounters after pair bonding, changes in DA receptor expression resulting from pair bonding may bias how aIC neurons “interpret” this DA signal to drive social valence-specific behaviors. aIC DRD1-expressing neurons project to DRD1-expressing neurons in the NAcSh and respond to social stimuli in mice [41], and NAcSh DRD1 receptors are required for the display of bond-specific behaviors in prairie voles [35, 36]. Thus, DA in the aIC may play an important and overlooked role upstream to modulate the pivotal role of NAc DA and its receptors in prairie vole pair bonding. We have previously observed changes in DA receptor expression in the aIC for prairie voles experience loss of a pair bonded partner [25]. These changes may serve an important role in social recognition and salience and valence assignment to specific social encounters or experiences. Proper integration of such socio-contextual information is a crucial first step when engaging in social encounters and forms the framework for establishing and maintaining a variety of social relationships.

## Acknowledgements

Special thank you to current and past members of the Smith Lab for the support and encouragement. We would also like to thank the KU Animal Care Unit staff for veterinary and husbandry care of the prairie vole colony at KU.

## Author Contributions

EV and AS conceptualized and designed all experiments; EV collected and analyzed all data with assistance from AT, KT, and KG; EV wrote the manuscript; AS and EV revised the paper for intellectual content following input and approval from all other authors.

## References

1. Lewis, A., Total umwelten create shared meaning the emergent properties of animal groups as a result of social signalling. Biosemiotics, 2020. 13(3): p. 431–441.

2. Lihoreau, M., et al., Context-dependent plasticity in social species: Feedback loops between individual and social environment. 2021, Frontiers Media SA. p. 645191.

3. Prior, N.H., E.J. Bentz, and A.G. Ophir, Reciprocal processes of sensory perception and social bonding: an integrated social℧sensory framework of social behavior. Genes, Brain and Behavior, 2022. 21(3): p. e12781.

4. O’Connell, L.A. and H.A. Hofmann, Evolution of a vertebrate social decision-making network. Science, 2012. 336(6085): p. 1154–1157.

5. Connell, L. and H. Hofmann, The Vertebrate Mesolimbic Reward System and Social Behavior Network: A Comparative Synthesis. 3639, 3599–3639. 2011.

6. Rogers-Carter, M.M. and J.P. Christianson, An insular view of the social decision-making network. Neuroscience & Biobehavioral Reviews, 2019. 103: p. 119–132.

7. Gehrlach, D.A., et al., A whole-brain connectivity map of mouse insular cortex. elife, 2020. 9: p. e55585.

8. Luchsinger, J.R., et al., Delineation of an insula-BNST circuit engaged by struggling behavior that regulates avoidance in mice. Nature Communications, 2021. 12(1): p. 3561.

9. Nicolas, C., et al., Linking emotional valence and anxiety in a mouse insula-amygdala circuit. Nature Communications, 2023. 14(1): p. 5073.

10. Duarte, I.C., et al., Ventral caudate and anterior insula recruitment during value estimation of passionate rewarding cues. Frontiers in Neuroscience, 2020. 14: p. 678.

11. Rogers-Carter, M.M., et al., Insular cortex mediates approach and avoidance responses to social affective stimuli. Nature neuroscience, 2018. 21(3): p. 404–414.

12. Kim, S.-H., et al., Anterior insula–associated social novelty recognition: pivotal roles of a local retinoic acid cascade and oxytocin signaling. American Journal of Psychiatry, 2023. 180(4): p. 305–317.

13. Girven, K.S., et al., Glutamatergic input from the insula to the ventral bed nucleus of the stria terminalis controls reward℧related behavior. Addiction biology, 2021. 26(3): p. e12961.

14. McGregor, M.S., C.V. Cosme, and R.T. LaLumiere, Insular cortex subregions have distinct roles in cued heroin seeking after extinction learning and prolonged withdrawal in rats. Neuropsychopharmacology, 2024: p. 1–10.

15. Starski, P., et al., Neural Activity in the Anterior Insula at Drinking Onset and Licking Relates to Compulsion-Like Alcohol Consumption. Journal of Neuroscience, 2024. 44(9).

16. Pribut, H.J., et al., Prior cocaine exposure increases firing to immediate reward while attenuating cue and context signals related to reward value in the insula. Journal of Neuroscience, 2021. 41(21): p. 4667–4677.

17. Bartels, A. and S. Zeki, The neural correlates of maternal and romantic love. Neuroimage, 2004. 21(3): p. 1155–1166.

18. Aragona, B.J. and Z. Wang, The prairie vole (Microtus ochrogaster): an animal model for behavioral neuroendocrine research on pair bonding. Ilar Journal, 2004. 45(1): p. 35–45.

19. Carter, C.S., A.C. Devries, and L.L. Getz, Physiological substrates of mammalian monogamy: the prairie vole model. Neuroscience & Biobehavioral Reviews, 1995. 19(2): p. 303–314.

20. Gossman, K.R., et al., Corticotropin-releasing factor and GABA in the ventral tegmental area modulate partner preference formation in male and female prairie voles (Microtus ochrogaster). Frontiers in Neuroscience, 2024. 18: p. 1430447.

21. Curtis, J.T. and Z. Wang, Ventral tegmental area involvement in pair bonding in male prairie voles. Physiology & behavior, 2005. 86(3): p. 338–346.

22. Aragona, B.J., et al., A critical role for nucleus accumbens dopamine in partner-preference formation in male prairie voles. Journal of Neuroscience, 2003. 23(8): p. 3483–3490.

23. Harbert, K.J., et al., How prior pair-bonding experience affects future bonding behavior in monogamous prairie voles. Hormones and behavior, 2020. 126: p. 104847.

24. Sherathiya, V.N., et al., GuPPy, a Python toolbox for the analysis of fiber photometry data. Scientific reports, 2021. 11(1): p. 24212.

25. Vitale, E.M., A. Kirckof, and A.S. Smith, Partner℧seeking and limbic dopamine system are enhanced following social loss in male prairie voles (Microtus ochrogaster). Genes, Brain and Behavior, 2023. 22(6): p. e12861.

26. de Kok, J.B., et al., Normalization of gene expression measurements in tumor tissues: comparison of 13 endogenous control genes. Laboratory investigation, 2005. 85(1): p. 154–159.

27. Tan, S.C., et al., Identification of valid housekeeping genes for quantitative RT-PCR analysis of cardiosphere-derived cells preconditioned under hypoxia or with prolyl-4-hydroxylase inhibitors. Molecular biology reports, 2012. 39: p. 4857–4867.

28. Schmittgen, T.D. and K.J. Livak, Analyzing real-time PCR data by the comparative CT method. Nature protocols, 2008. 3(6): p. 1101–1108.

29. Brusman, L.E., et al., Emergent intra℧pair sex differences and organized behavior in pair bonded prairie voles (Microtus ochrogaster). Genes, Brain and Behavior, 2022. 21(3): p. e12786.

30. Miura, I., et al., Encoding of social exploration by neural ensembles in the insular cortex. PLoS Biology, 2020. 18(9): p. e3000584.

31. Min, J.-Y., et al., The anterior insular cortex processes social recognition memory. Scientific Reports, 2023. 13(1): p. 10853.

32. Glangetas, C., et al., A population of Insula neurons encodes for social preference only after acute social isolation in mice. Nature Communications, 2024. 15(1): p. 7142.

33. Aragona, B.J. and Z. Wang, Dopamine regulation of social choice in a monogamous rodent species. Frontiers in behavioral neuroscience, 2009. 3: p. 649.

34. Gobrogge, K.L., et al., Anterior hypothalamic neural activation and neurochemical associations with aggression in pair℧bonded male prairie voles. Journal of Comparative Neurology, 2007. 502(6): p. 1109–1122.

35. Aragona, B.J., et al., Nucleus accumbens dopamine differentially mediates the formation and maintenance of monogamous pair bonds. Nature neuroscience, 2006. 9(1): p. 133–139.

36. Pierce, A.F., et al., Nucleus accumbens dopamine release reflects the selective nature of pair bonds. Current Biology, 2024. 34(3): p. 519-530. e5.

37. Ågmo, A., et al., Behavioral characteristics of pair bonding in the black tufted-ear marmoset (Callithrix penicillata). Behaviour, 2012. 149(3-4): p. 407.

38. Björnsdotter, M., et al., Somatotopic organization of gentle touch processing in the posterior insular cortex. Journal of Neuroscience, 2009. 29(29): p. 9314–9320.

39. Björnsdotter, M., I. Morrison, and H. Olausson, Feeling good: on the role of C fiber mediated touch in interoception. Experimental brain research, 2010. 207: p. 149–155.

40. Olausson, H., et al., Unmyelinated tactile afferents signal touch and project to insular cortex. Nature neuroscience, 2002. 5(9): p. 900–904.

41. Bellone, C., et al., Insular cortex to ventral striatum synapses encode valence of social interaction. 2022.

